# Ligand Identification in CryoEM and X-ray Maps Using Deep Learning

**DOI:** 10.1101/2024.08.27.610022

**Authors:** Jacek Karolczak, Anna Przybyłowska, Konrad Szewczyk, Witold Taisner, John M. Heumann, Michael H.B. Stowell, Michał Nowicki, Dariusz Brzezinski

**Affiliations:** Institute of Computing Science, Poznan University of Technology, Piotrowo 2, 60-965 Poznan, Poland; Department of Molecular, Cellular and Developmental Biology, University of Colorado Boulder, Boulder, CO 80309, USA; Institute of Robotics and Machine Intelligence, Poznan University of Technology, Piotrowo 3A, 60-965 Poznan, Poland

## Abstract

**Motivation:** Accurately identifying ligands plays a crucial role in the process of structure-guided drug design. Based on density maps from X-ray diffraction or cryogenic-sample electron microscopy (cryoEM), scientists verify whether small-molecule ligands bind to active sites of interest. However, the interpretation of density maps is challenging, and cognitive bias can sometimes mislead investigators into modeling fictitious compounds. Ligand identification can be aided by automatic methods, but existing approaches are available only for X-ray diffraction and are based on iterative fitting or feature-engineered machine learning rather than end-to-end deep learning.

**Results:** Here, we propose to identify ligands using a deep learning approach that treats density maps as 3D point clouds. We show that the proposed model is on par with existing machine learning methods for X-ray crystallography while also being applicable to cryoEM density maps. Our study demonstrates that electron density map fragments can aid the training of models that can later be applied to cryoEM structures but also highlights challenges associated with the standardization of electron microscopy maps and the quality assessment of cryoEM ligands.

**Availability:** Code and model weights are available on GitHub at https://github.com/jkarolczak/ligands-classification. Datasets used for training and testing are hosted at Zenodo: 10.5281/zenodo.10908325. An accompanying ChimeraX bundle is available at https://github.com/wtaisner/chimerax-ligand-recognizer.

**Supplementary information:** Supplementary data are available at *Bioinformatics* online.

## 1 Introduction

X-ray crystallography and cryogenic-sample electron microscopy (cryoEM) are currently the most popular techniques for determining the 3D structures of proteins and nucleic acids. Many such structures contain small-molecule ligands that can illuminate the function of the macromolecule they bind to. Therefore, the correct identification of ligands is often a vital part of structure-guided drug design. However, ligands are usually modeled manually by chemists or biologists analyzing 3D density maps. This process is time-consuming and prone to human error, particularly for structures with low resolution or local disorder. As a result, several studies have reported questionable assignments of ligands to density fragments (Deller and Rupp, 2015; Wlodawer *et al*., 2018).

Automatic ligand identification and fitting methods have been proposed to aid structural biologists in modeling small molecules. When the ligand to be modeled is known, iterative fitting procedures based on core atom recognition, followed by iterative element addition and optimization, can be used (Terwilliger *et al*., 2006; Zwart *et al*., 2004; Evrard *et al*., 2007; Muenks *et al*., 2023). These methods can be adapted to identify unknown ligands by fitting moieties from a predefined list of candidates (Terwilliger *et al*., 2007), however, such an approach can be slow as it requires trial fitting of all candidate ligands. Alternatives to iterative fitting are based on statistical descriptions of 3D density map fragments and machine learning (Aishima *et al*., 2005; Gunasekaran *et al*., 2009; Carolan and Lamzin, 2014; Kowiel *et al*., 2019). Notably, the recently proposed CheckMyBlob algorithm and web server (Kowiel *et al*., 2019; Brzezinski *et al*., 2021) are faster and more accurate than iterative fitting approaches. However, all the existing ligand prediction approaches are applicable only to X-ray structures and are based on manually engineered features rather than end-to-end deep learning from density maps.

Deep learning is already being applied to structural biology. Convolutional neural networks and vision transformers have been used to analyze X-ray diffraction images (Czyzewski *et al*., 2021; Banko *et al*., 2021) and cryoEM micrographs (Bepler *et al*., 2019; Dhakal *et al*., 2024). Deep learning approaches have also been developed for monitoring the crystallization process (Ito *et al*., 2019; Matinyan *et al*., 2024), protein structure prediction (Jumper *et al*., 2021; Baek *et al*., 2021), structure determination (Pan *et al*., 2023; Li *et al*., 2024), cryoEM map improvement (Sanchez-Garcia *et al*., 2021; He *et al*., 2023), side-chain conformation prediction (Misiura *et al*., 2022), protein classification (Weiler *et al*., 2018), and model building (Jamali *et al*., 2024). However, all the methods above focus on raw experimental data or macromolecules rather than small-molecule ligands. Furthermore, it is well-recognized that ligand identification and proper fitting are ongoing challenges in the cryoEM field (Lawson *et al*., 2024).

Even though deep learning has not been used to identify ligands based on density map fragments, several deep learning classification methods for 3D objects already exist. Data regarding 3D objects is often gathered through LiDAR scanning and stored as point clouds. Such data can be analyzed using point cloud neural network architectures called pointnets (Charles *et al*., 2017; Qi *et al*., 2017). However, everyday objects are usually scanned in one (‘upright’) position, and regular pointnets would not work well with ligands, which do not have a predefined orientation. To solve point cloud orientation problems, rotation-invariant pointnets have been proposed, such as the recent RiConv and RiConv++ algorithms (Zhang *et al*., 2019, 2022). Some form of translation and rotation invariance is also needed in LiDAR-based place recognition, where algorithms such as MinkLoc3D (Komorowski, 2021, 2022; Zywanowski *et al*., 2022) and TransLoc3D (Xu *et al*., 2023) have shown promising results. Therefore, there are several deep learning architectures that could be applied to 3D ligand shape recognition from experimental density maps.

In this paper, we present an end-to-end deep learning approach to identifying ligands in 3D density map fragments. Herein, we compare several density map sampling strategies and test rotation-invariant pointnets and sparse convolutional networks. The deep learning models are trained on electron density maps from X-ray crystallography, but in contrast to existing methods, they can also be applied to Coulomb potential maps from cryoEM. Experiments assessing model performance on 208,896 X-ray crystallography ligands show that the proposed approach has similar accuracy to existing methods while improving in terms of top-10 accuracy. Experiments involving 34,671 cryoEM ligands show that this performance can be translated to cryoEM and mixed sets of ligands. We make the proposed model available as a ChimeraX bundle to facilitate adoption. Finally, we discuss the limitations of current cryoEM map processing and quality assessment procedures.

## 2 Materials and methods

### 2.1 Data collection and processing

#### 2.1.1 X-ray crystallography data

For training the ligand classification model, we used the same source data and map processing methods as described by Brzezinski *et al*. (2021), consisting of structures downloaded from the Protein Data Bank (PDB) (Berman, 2000) as of 19 January 2020. Using the downloaded structures, we extracted 957,855 ligand blobs. The ligands were identified by positive electron density peaks within the F_o_-F_c_ map limited by the 2.8σ isosurface computed with a 0.2 Å grid. Each detected ligand was saved as a 3D voxel grid with 0.2 Å spacing. Suspicious deposits and ligands were eliminated according to the following quality criteria: resolution > 4.0 Å, RSCC < 0.6, real space Z_obs_ (RSZO) < 1.0, real space Z_diff_ (RSZD) ≥ 6.0, R factor > 0.3, or occupancy < 0.3. The resulting dataset consisted of 696,887 blobs with initial labels assigned based on residue identifiers in the PDB deposits. Since several ligands are indistinguishable by electron density alone, we followed the procedure used by the CheckMyBlob server and clustered ligands into ligand groups based on the number of atoms, number of rings, connectivity, chirality, and the atomic numbers of corresponding atoms (Brzezinski *et al*., 2021). Clustering was performed using RDKit (http://www.rdkit.org) based on the SMILES and InChI descriptors provided by the PDB. We then limited the number of labels used for classifier training to ligand groups with at least 100 blob instances. All the ligands not in those 218 groups were labeled as a separate class called rare, creating a total of 219 ligand groups. Further details concerning the blob extraction and ligand label assignment methods can be found in (Kowiel *et al*., 2019; Brzezinski *et al*., 2021).

#### 2.1.2 CryoEM data

For testing the final model on cryoEM data, we downloaded 6,103 EM-derived CIF models of proteins containing ligands. We used structures with reported resolutions of 4.0 Å or better from the PDB as of 30 November 2023, along with the corresponding EM density maps from the Electron Microscopy Data Bank (EMDB) (The wwPDB Consortium *et al*., 2024). Complete map-vs-original-model Q-scores were computed using the Chimera mapq plug-in (Pintilie *et al*., 2020), and individual ligand Q-scores were extracted. Ligand-free models were generated by deleting all HETATM entries, and difference maps between these ligand-free models and the EMDB maps were created with the Phenix command phenix.real_space_diff_map (Liebschner *et al*., 2019). Blobs corresponding to individual ligands were extracted, and those with Q-scores of 0.6 or better and volume greater than V = 2.14 Å^3^ were processed. The filtering resulted in a dataset of 34,671 ligand blobs labeled using the same 219 groups that were assigned to X-ray ligands. Each extracted cryoEM ligand was resampled and represented as a 3D voxel grid with 0.2 Å spacing, the same format that was used for X-ray crystallography blobs.

#### 2.1.3 CryoEM map transformation

The processing of X-ray diffraction data has been standardized over the years, therefore, the representation of ligands by thresholding the F_o_-F_c_ map at 2.8σ produces comparable results for all PDB deposits (Supplementary Fig. S1A). Our goal was to produce similar maps for cryoEM ligands. However, EM Coulomb potential maps are still being processed differently by different labs. In particular, the varying resolution of different map fragments and various forms of sharpening, sometimes position-dependent, make it impossible to select a single threshold for the density analysis of all cryoEM maps. Although thresholding by false discovery rate (FDR) can successfully reduce background noise and help visually interpret macromolecules (Beckers et al., 2019), FDR thresholding produces binary density maps that are often noisy at the ligand level. As a result, our attempt to use FDR thresholding to create 3D ligand representations resulted in voxel grids that had discontinuities and differed significantly from the grids obtained from X-ray crystallography. Therefore, for the purposes of this study, we have developed a custom cryoEM map normalization and thresholding method.

The proposed ligand map normalization method consists of three steps: *reducing zero inflation, quantile thresholding*, and *voxel value normalization*. Many cryoEM maps have their volume masked during refinement, zeroing or solvent-flattening voxel values outside the selected (particle) region. Depending on the map box size, which is somewhat arbitrary and macromolecular molecule size dependent, masking introduces varying fractions of low-value voxels in the map. As a result, the distribution of map voxel values of a cryoEM map usually contains a spike around zero (Fig. 1A, Supplementary Fig. S1B), where the size of the spike depends on the relative size of the mask and box (Afonine *et al*., 2018). Electron density difference maps from crystallography do not have this variability and have close to normal distributions (Supplementary Fig. S1A). This is partly because the box always spans a full unit cell, without the need for masking and without variable padding around the region of high density. To reduce the effect of zero-inflation introduced by masking and arbitrary box size selection, when determining the density cutoff threshold, we analyze the histogram of the entire cryoEM map (Fig. 1A) and ignore values +/-0.5 standard deviation around the median of voxel values (Fig. 1B). Next, since the resulting map density histogram is usually still not normally distributed, we use quantile thresholding instead of standard deviation thresholding. We have selected the quantile corresponding to 2.8σ in a normal distribution, which translates to a cumulative distribution value of approximately 0.9974. Finally, after performing the density cutoff on the difference map and zeroing voxel values below the selected threshold, the ligand density voxel values are normalized to resemble electron density values from X-ray crystallography. For this purpose, we multiplicatively rescale voxel values so that the lowest non-zero value is equal to the average lowest non-zero value of X-ray ligands for the given resolution. The 3D voxel grids of X-ray and cryoEM ligand blobs (in compressed numpy array format) are hosted at Zenodo: 10.5281/zenodo.10908325.

**Fig. 1.**
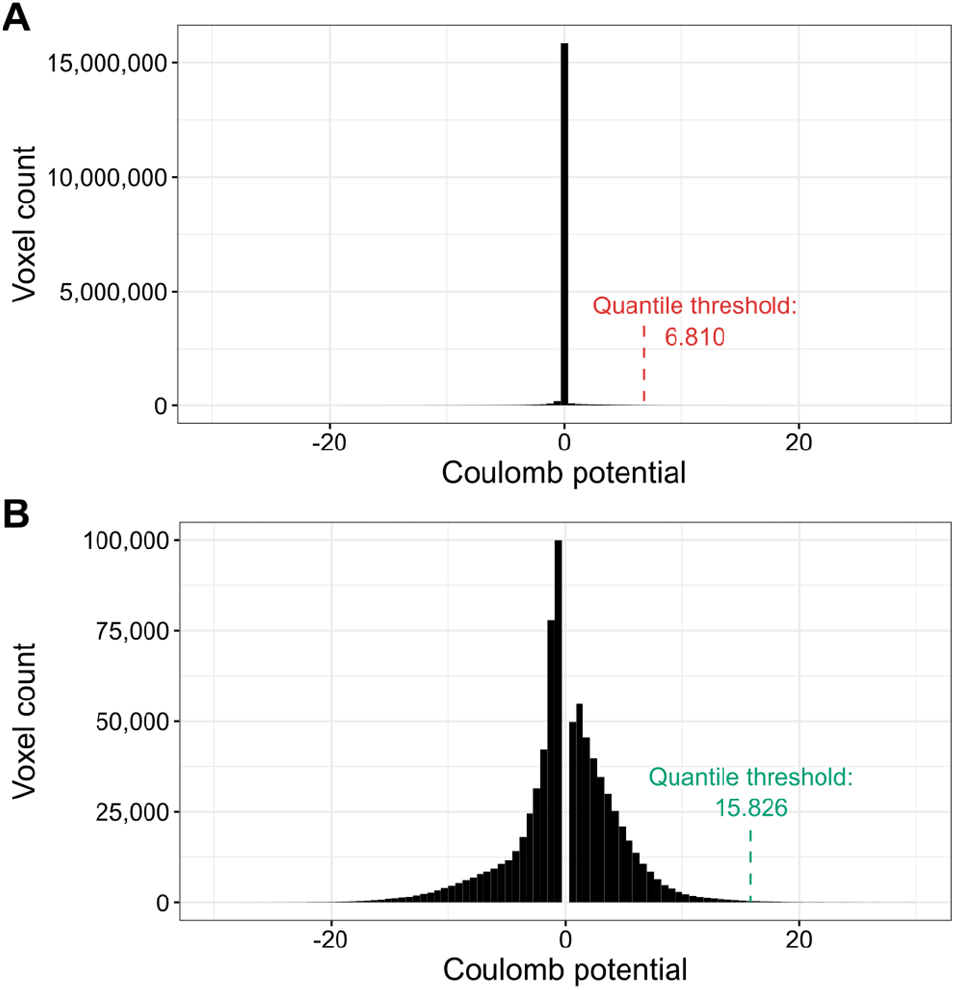
Illustration of the density thresholding used for cryoEM maps. (**A**) The default voxel value distribution for PDB deposit 7SMR. The distribution is zero-inflated (almost 15 million voxels with values close to zero). (**B**) Voxel value distribution of 7SMR after the removal of values +/-0.5 standard deviation around the median. The resulting distribution is much closer to normal and the quantile threshold is over two times higher.

**Fig. 2.**
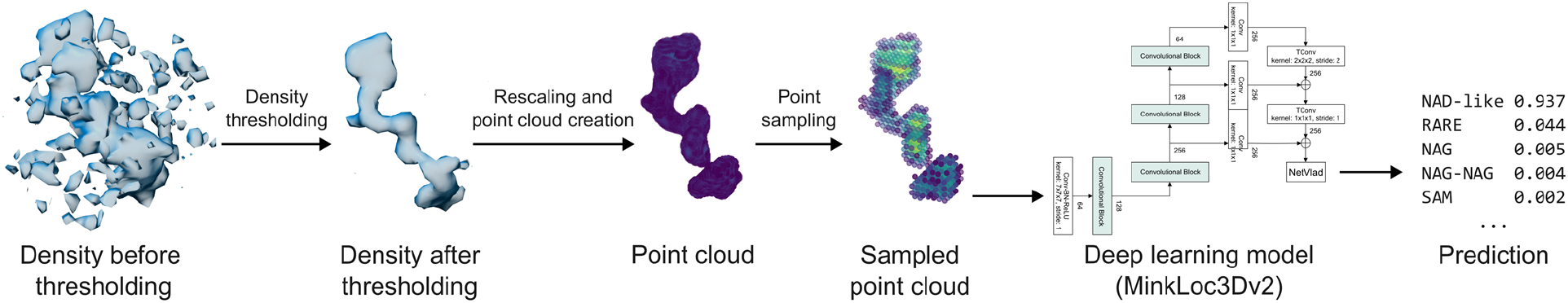
Schematic representation of the proposed ligand recognition pipeline. Example presenting the processing of a nicotinamide adenine dinucleotide ligand from 2.10 Å cryoEM PDB deposit 8fuz (residue A 602). An initially selected fragment of a difference map (represented here at a 2.8σ threshold) is given a new density threshold based on reducing zero inflation and quantile thresholding. The map fragment is then resampled into a 0.2 Å grid, voxel values are rescaled, and the grid is transformed into a point cloud. From the point cloud, 2000 points are selected (using uniform maximum point selection) and fed into a deep learning model that outputs a list of ligand group scores.

### 2.2 Design and implementation of deep learning models

Based on an analysis of literature concerning position-invariant 3D object recognition, we selected three deep learning architectures as end-to-end ligand recognition models: RiConv++ (Zhang *et al*., 2022), TransLoc3D (Xu *et al*., 2023), and MinkLoc3Dv2 (Komorowski, 2022). In the following subsections, we describe how the selected architectures were adapted to the problem of ligand recognition based on 3D voxel grids. An overview of the architectures is presented in Supplementary Figure S2.

#### RiConv++

RiConv++ (Zhang *et al*., 2022) is a deep learning architecture developed to enhance the rotation-invariant convolution (RIConv, Zhang *et al*., 2019) model for 3D point cloud data processing. The main improvements in RiConv++ include introducing the use of Local Reference Axis (LRA) or normal vectors to calculate more stable and descriptive Informative Rotation-Invariant Features (IRIF). RiConv++ projects local points onto a plane perpendicular to the LRA, using spatial relations to derive a comprehensive description of point neighborhoods. Our adaptation of the RiConv++ architecture incorporated five layers of RIConv++ operators, each followed by batch normalization and a rectified linear unit (ReLU) layer. The estimation of the probabilities of the 219 ligand groups was performed by two fully connected layers followed by a softmax layer.

#### TransLoc3D

TransLoc3D (Xu *et al*., 2023) is a recent model for place recognition in 3D point cloud environments. The architecture of TransLoc3D incorporates several advanced components, including 3D Sparse Convolution, Adaptive Receptive Field, External Transformer, and NetVLAD. The 3D Sparse Convolution module aggregates local geometric information efficiently thanks to sparse convolutional layers. The Adaptive Receptive Field mechanism captures structure information from different neighborhood sizes through parallel branches with varying receptive fields, refined further by Efficient Channel Attention (ECA). The model also utilizes an External Transformer to maintain contextual information across nearby and distant points, implementing efficient attention mechanisms. Finally, NetVLAD aggregates local features into a global descriptor by summing residuals between local features and cluster centers, thus enhancing the model’s ability to produce a robust global representation of ligands.

#### MinkLoc3Dv2

MinkLoc3Dv2 (Komorowski, 2022) is an enhanced version of the MinkLoc3D architecture (Komorowski, 2021) designed for place recognition in 3D environments. This model extends the depth and breadth of the original by increasing the number of convolutional and transposed convolutional blocks, as well as the number of channels within the network. Each convolution block is further enhanced with Efficient Channel Attention (ECA) modules to improve local cross-channel interactions, which were absent in its predecessor. Our implementation is a U-Net-like architecture inspired by the Feature Pyramid Network concept, employing a bottom-up pathway composed of three convolution blocks with increasing receptive fields to produce feature maps and a top-down pathway with transposed convolutions to add these features back to the network at corresponding levels. We modified the final global descriptor, which originally consisted of Generalised-Mean (GeM) pooling, to use NetVLAD (Arandjelovic *et al*., 2016) to classify ligands.

#### Point sampling and hyperparameter tuning

Each ligand type has a different size and shape, resulting in differently-sized voxel grids. For instance, a phosphate ion (PO4) consists of 5 non-hydrogen atoms, whereas heme (HEM) has 43. Therefore, estimating the memory required for processing a voxel-based point cloud representation of a ligand is difficult. As a result, the training process on point clouds consisting of all ligand voxels would be highly unstable, with big ligands potentially interrupting the process by causing out-of-memory errors. Therefore, we tested several transformations that aimed at limiting the maximal number of points used to represent a ligand while retaining as much information about its shape as possible. In other words, our goal was to use only selected voxels of each ligand so that its size would not exceed a defined threshold.

To limit the processed points to a predefined number *max*_*p*_, we investigated four sampling strategies: *random, uniform, surface*, and *clustering* (Supplementary Fig. S3). Random sampling randomly selects non-zero points from the point cloud until the desired number of points is met. Uniform sampling divides the point cloud into a grid of equally sized meta-voxels in such a way that the number of meta-voxels with non-zero points is ≤ *max*_*p*_. The number of meta-voxels is obtained iteratively by first creating one meta-voxel comprising the entire original blob voxel grid and then dividing it into smaller meta-voxels. For this, we define a divisor *N* that cuts the initial grid into *N*^3^ meta-voxels (each face of the grid is divided by *N*). For instance, a point cloud with initial grid dimensions (20, 16, 10) using *N*=2 will be divided into 8 blocks, each of shape (10, 8, 5). Non-divisible dimensions are zero-padded. *N* is iteratively increased by 1 to achieve the maximal number of meta-voxels with non-zero points as close to *max*_*p*_ as possible. Finally, from each meta-voxel we select one point— the one with maximum density. Importantly, the sampled point retains its coordinates and density value at the original scale. The surface sampling strategy uniformly samples points from the outer shell of a ligand, whereas the clustering sampling strategy performs *k*-means clustering (*k = max*_*p*_) and uses centroids as the final points.

Hyperparameter tuning experiments (Supplementary Table S1) have shown that the best sampling strategy was uniform sampling with selecting the maximum point within a meta-voxel, and *max*_*p*_ was set to 2000 points. Apart from deciding on the point sampling strategy, results on a validation portion of the training set were used to determine the learning rate, number of batches in gradient accumulation, ligand under-/over-sampling, and number of epochs.

### 2.3 Machine learning experimental setup

To evaluate the recognition rate of the trained deep learning models (RiConv++, TansLoc3D, MinkLoc3Dv2) and CheckMyBlob (Kowiel *et al*., 2019), the collected data were divided to evaluate selected models in three settings: *X-ray, cryoEM*, and *mixed* ligand type test sets. For the X-ray evaluation, we used a stratified sample of 70% (486,991) of the X-ray ligands as training data and held out the remaining 30% (208,896) for testing. Stratification was particularly important in this study, as the collected data has a strongly skewed ligand type distribution (Kowiel *et al*., 2019), and purely random, non-stratified holdout would produce unreliable error estimates. Moreover, ligands belonging to the same PDB deposit were all assigned either to the training set or testing set to avoid training and testing data from the same PDB structure. For the purposes of hyperparameter tuning, 25% of the training set was used as an independent validation set. We note that the CheckMyBlob model was trained and tested on exactly the same ligands as the deep learning models. For cryoEM evaluation, the 34,671 cryoEM ligands were divided into three folds and the models were evaluated using stratified 3-fold cross-validation. For the mixed training regime, the model was trained on the 70% stratified X-ray sample used for X-ray evaluation and two cryoEM folds from the cryoEM evaluation and then tested on the X-ray holdout and one cryoEM fold.

The classifiers were evaluated using the following metrics: classification accuracy, top-10 accuracy, mean correct prediction rank, Brier score, and macro-averaged recall. Classification accuracy is the proportion of correctly recognized ligands among all testing examples. Top-10 accuracy is the proportion of cases where the correct ligand was among the ten highest-ranked hits in the classifier’s prediction. Mean correct prediction rank is the average position of the correct prediction on the list of ligand probabilities. Brier score measures the squared probability estimation error for the correct class, whereas macro-averaged recall is the (unweighted) arithmetic mean of the recognition rates for each of the 219 ligand groups. All the selected measures are commonly used in machine learning to evaluate classifiers on datasets with skewed class distributions (Japkowicz and Shah, 2011). The code for the experiments is available on GitHub: https://github.com/jkarolczak/ligands-classification.

## 3 Results

### 3.1 Experimental comparison of end-to-end deep learning and feature-based ligand recognition

The predictive performance of the proposed deep learning models and the feature-based CheckMyBlob algorithm on X-ray and cryoEM ligands is summarized in Table 1.

**Table 1.** Prediction results on test datasets. The best values for each evaluation measure are shown in bold, and the runner-up is underlined.

| Dataset | Algorithm | Acc. | Top-10 Acc. | Pred. rank | Brier score | Macro recall |
| --- | --- | --- | --- | --- | --- | --- |
| X-ray | CheckMyBlob | **0.672** | <u>0.936</u> | <u>3.716</u> | **0.331** | **0.350** |
|  | RiConv++ | 0.542 | 0.920 | 4.410 | 0.478 | 0.104 |
|  | TransLoc3D | 0.606 | 0.930 | 4.196 | 0.399 | 0.208 |
|  | MinkLoc3Dv2 | <u>0.660</u> | **0.946** | **3.627** | <u>0.344</u> | <u>0.325</u> |
| CryoEM | MinkLoc3Dv2<sup>X</sup> | 0.344 | 0.851 | 8.323 | 0.641 | 0.044 |
|  | MinkLoc3Dv2<sup>C</sup> | <u>0.639</u> | <u>0.930</u> | <u>4.776</u> | <u>0.385</u> | **0.168** |
|  | MinkLoc3Dv2<sup>X+C</sup> | **0.691** | **0.974** | **2.589** | **0.280** | <u>0.157</u> |
| Mixed | MinkLoc3Dv2 | **0.658** | **0.946** | **3.723** | **0.342** | **0.330** |

Acc.=Accuracy, Pred. rank=Correct prediction rank, <sup>X</sup>=X-ray training data, <sup>C</sup>=cryoEM training data, <sup>X+C</sup>=X-ray and cryoEM training data

Looking at the results, one can notice that the TransLoc3D and RiConv++ models generally underperformed compared to MinkLoc3Dv2 and CheckMyBlob on the X-ray data. MinkLoc3Dv2 achieved the highest top-10 accuracy (0.946) and the best mean correct prediction rank (3.627), suggesting its strong ability to place the correct ligand among the top candidates. This ranking consistency is crucial for applications where reviewing top suggestions rather than a single prediction is viable. The overall accuracy of MinkLoc3Dv2 (0.660) was only slightly lower than CheckMyBlob (0.672), indicating comparable performance in exact ligand identification. Interestingly, CheckMyBlob maintained the highest macro recall (0.350) for X-ray data, indicating better accuracy across all ligand classes, including rare ones.

The performance on cryoEM data was tested only with MinkLoc3Dv2, as CheckMyBlob cannot be applied to Coulomb potential maps because it uses electron count and contour level dependent features, and TransLoc3D and RiConv++ underperformed. Due to the relatively small number of training examples for cryoEM, we investigated three training schemes: training only on X-ray data (MinkLoc3Dv2^X^), training only on cryoEM data (MinkLoc3Dv2^C^), training on a mixture of X-ray and cryoEM ligands (MinkLoc3Dv2^X+C^). In all three cases, the models were evaluated on the same set of (only cryoEM) ligands. Comparing the predictive performance of these three training scenarios, one can see a clear benefit of combining X-ray and cryoEM data. Even though using X-ray data alone gives relatively poor results, its addition to cryoEM ligands substantially boosts accuracy, top-10 accuracy, and mean correct prediction rank. To verify whether this performance holds on a test set involving both X-ray and cryoEM ligands, we also tested MinkLoc3Dv2 trained on mixed types of ligands. As can be seen in Table 1, the results in the mixed training scenario are similar to those on X-ray and cryoEM.

Figure 3 provides further insights into the models’ performance across different map resolutions, ligand quality, and ligand sizes. For X-ray data, the mean rank of correct ligand predictions for all models improves as resolution increases (Fig. 3A) and as the real-space correlation coefficient (RSCC) increases (Fig. 3B). This trend is expected as higher resolution and RSCC values typically indicate better quality data. There is no clear correlation between ligand size and model performance (Fig. 3C). Interestingly, for cryoEM ligands, the performance of MinkLoc3Dv2 does not seem to depend on resolution, ligand Q-score quality or ligand size (Fig. 3D–F). What is visible, however, is the consistently improved predictive performance of the model trained on X-ray and cryoEM ligands (MinkLoc3Dv2^X+C^) compared to models trained on only ligands from one type of map. Finally, the results for the mixed training and testing data resemble those for X-ray data, with slightly improved performance relative to that on X-ray data for lower resolution, poorer quality, and larger ligands (Fig. 3G–I).

**Fig. 3.**
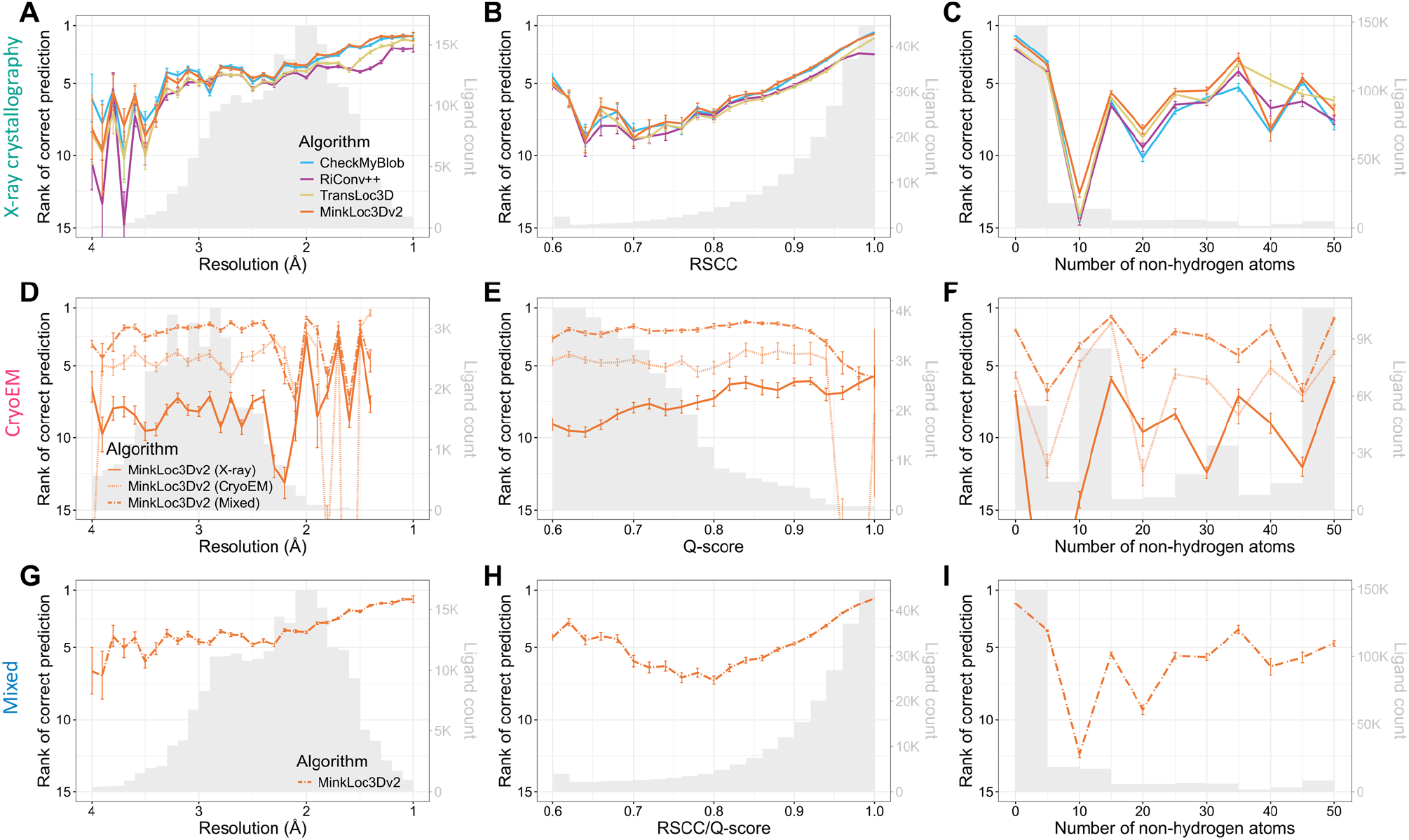
Predictive performance of the analyzed machine learning models for X-ray (top panels), cryoEM (middle panels), and mixed (bottom panels) test sets of ligands. Each plot shows the mean ranks of the correct ligand (with error bars showing the standard error) versus (**A, D, G**) map resolution, (**B, E, H**) ligand fit quality (RSCC for X-ray, Q-score for cryoEM), and (**C, F, I**) size (number of non-hydrogen atoms) of the ligand. The corresponding ligand distributions are depicted as gray histograms, with ligand counts shown on the right y-axis.

We also investigated model performance plotted jointly against resolution and ligand quality (Supplementary Fig. S4) and resolution and ligand size (Supplementary Fig. S5). It can be noticed that the deep learning models obtain higher mean prediction ranks than CheckMyBlob for lower resolution, poorer quality, and larger ligands. Indeed, when one investigates the performance of the models per ligand group, it seems that CheckMyBlob achieves better accuracy, mostly for popular ligands (Supplementary File 1, Fig. S6-S8). This results in the overall high performance of CheckMyBlob on the X-ray dataset due to the high imbalance between different ligand groups (Supplementary Fig. S9). Finally, the models perform similarly well regardless of the size of the initial ligand point cloud (Supplementary Fig. S10).

It is also worth pointing out the differences in the measured quality of X-ray and cryoEM ligands. As can be noticed by looking at the gray histograms in Fig. 3B and 3E, there are far fewer high-quality cryoEM ligands compared to X-ray ligands. Whereas most X-ray diffraction ligands have RSCC between 0.9 and 1.0, the majority of cryoEM ligands have Q-scores below 0.8. Of course, RSCC and Q-score are different metrics and, therefore, should not be compared directly (although their intuitions and formulas are similar (Pintilie *et al*., 2020)). Nevertheless, the shapes of the RSCC and Q-score distributions show a clear gap between the quality of X-ray and cryoEM ligands. This may partly be due to the fact that Q-scores seem to be correlated with resolution (Supplementary Fig S4B), and resolution is generally lower for cryoEM.

We have also verified that all the tested models are generally well-calibrated, i.e., their prediction probabilities correspond linearly with their recognition rates (Supplementary Fig. S11). The only exception was the MinkLoc3Dv2^X^ model trained on X-ray ligands and tested on cryoEM ligands, highlighting the inconsistencies between X-ray and cryoEM map contouring thresholds.

In addition to analyzing the predictive performance, we also compared the running times of selected models. On a set of 100 randomly chosen ligands, MinkLoc3Dv2 needed 0.103 seconds to process a single ligand, whereas CheckMyBlob needed 3.991 seconds (measurements for both algorithms were done on a single core of an Intel i7-7700HQ CPU). This significant difference stems from the fact that CheckMyBlob spends compute time sequentially calculating all the blob descriptors (features), whereas MinkLoc3Dv2 is an end-to-end approach that encodes an internal representation of a blob during inference. The prediction speed of the proposed deep learning approach may be important in applications where predictions are performed at high rates, e.g., as part of fragment screening campaigns (Pearce *et al*., 2017).

### 3.2 Analysis of predicted ligands

To additionally validate the performance of the proposed MinkLoc3Dv2 model, we inspected selected predictions on X-ray and cryoEM ligands (Fig. 4). We looked at X-ray diffraction ligands from PDB entries: 4iun, which was analyzed in previous ligand prediction studies (Carolan and Lamzin, 2014; Kowiel *et al*., 2019); 3nw4, which illustrates the recognition of buffer components; 6nau and 2acw, which show situations where MinkLoc3Dv2 correctly predicted the ligand and CheckMyBlob did not; and 3i0l, which highlights a situation where CheckMyBlob predicted correctly and MinkLoc3Dv2 focused on only part of the density. We have also inspected cryoEM predictions by looking at PDB deposits 7qh2, 6zku, 8fuz, and 8hdp, which show correct identifications of small, medium, and large ligands at different resolutions, as well as entry 7jro, which highlights the problem of map thresholding.

**Fig. 4.**
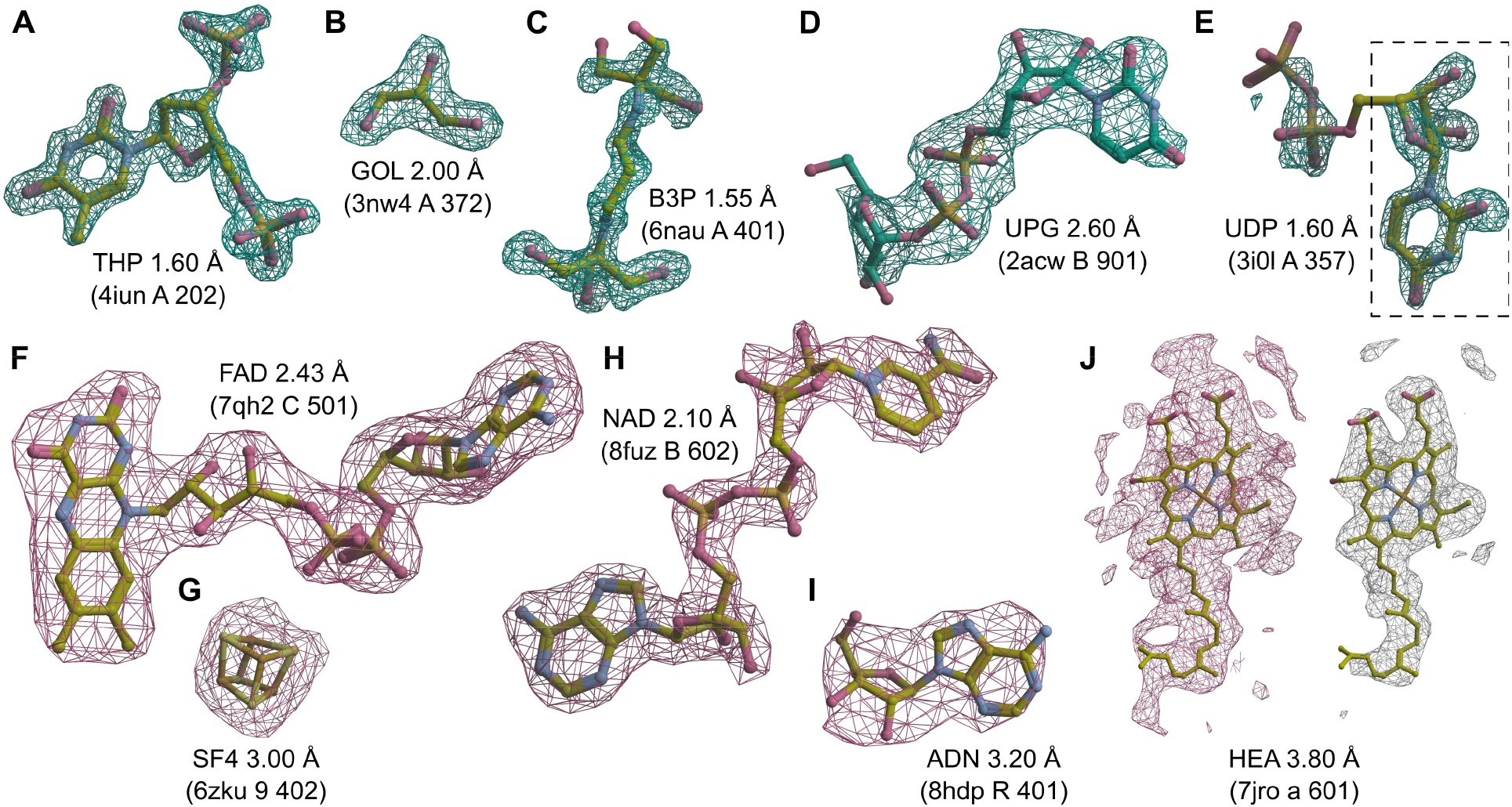
Examples of ligand identification using the proposed MinkLoc3Dv2 model. (**A–D**) Examples of correctly predicted X-ray ligands. (**E**) Uridine-5’-diphosphate (UDP) misclassified as uridine (URI, black dashed frame). (**F–I**) Examples of correctly predicted cryoEM ligands. (**J**) Heme A (HEM) misclassified as a rare ligand due to incorrect density thresholding. Each ligand is labeled by its Chemical Component Dictionary ID, structure resolution, and (in parentheses) the PDB ID, chain, and residue number. X-ray diffraction ligands shown in green mesh based on F_o_-F_c_ maps contoured at 2.8σ calculated after removal of solvent and other small molecules (including the ligand) from the model. CryoEM ligands depicted in pink mesh based on difference maps contoured according to the proposed automatic density thresholding method (13.642, 3.385, 17.997, 7.850, and 5.613 V for panels F–J, respectively). The white mesh in panel J shows a manually selected contour threshold of 11.000 V. Atomic coordinates were taken from the PDB deposits.

MinkLoc3Dv2 was able to identify large and medium-sized ligands such as thymidine-3’,5’-diphosphate (Fig. 4A), 2-[3-(2-hydroxy-1,1-dihydroxymethyl-ethylamino)-propylamino]-2-hydroxymethyl-propane-1,3-diol (Fig. 4C), and uridine-5’-diphosphate-glucose (Fig. 4D). Moreover, the model recognizes common buffer or cryo-protectant components, such as glycerol (Fig. 4B). MinkLoc3Dv2 was able to provide correct predictions when CheckMyBlob was not in cases with missing electron density at terminal atoms (Fig. 4C and 4D). On the other hand, MinkLoc3Dv2 had trouble identifying the correct ligand when the electron density was poorly defined, resulting in discontinuous blobs. An example of such a situation can be observed in Fig. 4E, where the deep learning model predicted uridine (URI) instead of uridine-5’-diphosphate (UDP). This misidentification results from the fact that the highest electron density peak (black dashed frame in Fig. 4E) corresponds to URI, a ‘component’ of UDP. Nevertheless, MinkLoc3Dv2 and CheckMyBlob agreed in most cases we inspected, and when one of the models misclassified, the correct ligand was usually among the top 10 predictions.

A distinctive property of MinkLoc3Dv2 is that it can be used to predict not only X-ray ligands but also cryoEM ligands. MinkLoc3Dv2 was able to identify ligands of various sizes, such as flavin adenine dinucleotide (Fig. 4F), an iron-sulfur cluster (Fig. 4G), nicotinamide adenine dinucleotide (Fig. 4H), and adenosine (Fig. 4I). In all of these cases, the proposed automatic map thresholding method worked well. However, there were also cases where the automatically determined contour level was suboptimal. In the case of heme A depicted in Fig. 4J, the automatic threshold of 5.613 V (pink mesh) was too low, resulting in an incorrect prediction. The ligand-to-map fit at a manually set 11.000 V contour level (white mesh) indicates that the prediction could have been more accurate at a different threshold. This shows that inconsistencies between cryoEM maps, caused by different Coulomb potential ranges, B-factor compensation or other sharpening, and varying resolution for different map fragments, are possibly the main obstacles for automatic data processing and machine learning on cryoEM ligands.

## 4 Discussion

The application of deep learning to ligand identification in X-ray and cryoEM maps reveals both promising results and significant challenges. Our experiments demonstrate that deep learning approaches can achieve comparable accuracy and better top-10 accuracy than existing feature-based methods for X-ray crystallography data while also being applicable to cryoEM structures. To the best of our knowledge, this is the first deep learning approach to recognize X-ray ligands and the first approach of any type to automatically identify ligands in cryoEM. To facilitate the adoption of our method by biologists, we have developed a plugin (bundle) that makes it possible to predict ligands within ChimeraX.

Even though the proposed MinkLoc3Dv2 model offers similar predictive performance for X-ray and cryoEM ligands, our study highlights fundamental differences between these two experimental. X-ray crystallography and cryoEM produce different types of maps—electron density maps and Coulomb potential maps, respectively. While both aim to reveal the 3D structure of macromolecules, they differ in several key aspects. X-ray maps represent the distribution of electrons, while cryoEM maps show the electrostatic potential, which is influenced by both electrons and nuclei (Wang, 2017). However, X-ray maps and cryoEM maps are related (Marques *et al*., 2019). Indeed, it has been shown that (until quite high resolution) the electron density and potential are similar in shape (Mitsuoka *et al*., 1999). This relation allowed us to successfully predict ligands in cryoEM maps using X-ray maps as part of the training data.

A main challenge in applying deep learning models trained on X-ray data to cryoEM comes from the lack of standardization in cryoEM map processing. Unlike X-ray crystallography, where data processing has been refined over decades, cryoEM map generation and post-processing methods can vary significantly between research groups (Rosenthal and Henderson, 2003). This makes it difficult to establish consistent thresholds for map interpretation and comparison. Our attempts to use false discovery rate (FDR) thresholding, while successful for overall macromolecule visualization (Beckers *et al*., 2019), proved inadequate for extracting consistent ligand representations. The custom normalization and thresholding method we developed for this study is a step towards addressing this issue, but further work is needed to establish community-wide standards for cryoEM map processing. Map thresholding strongly affects blob extraction, which is a key component that can affect the performance of the analyzed algorithms in practical applications.

The architecture of deep learning models for ligand recognition is also a topic open for future work. The fact that the MinkLoc3Dv2 model, which is not designed to support complete rotation invariance, worked much better than the fully rotationally invariant RiConv++ is a somewhat surprising result. Therefore, further research in the field of 3D computer vision is needed to fully understand the key neural network components needed for accurate and robust ligand classification.

The validation of ligands in cryoEM structures presents another significant challenge. While robust validation metrics and procedures exist for X-ray structures (Smart *et al*., 2018), equivalent measures for cryoEM are still evolving. The Q-score metric we used in this study provides valuable information about local map quality (Pintilie et al., 2020) and has been recently recognized by the community as a useful tool for macromolecule and ligand validation (Burley *et al*., 2022; Lawson *et al*., 2024). Nevertheless, our analysis revealed a clear gap in the quality distribution between X-ray and cryoEM ligands, with most cryoEM ligands having Q-scores far below 0.8. The weak correlation between model performance and Q-score suggests a need for additional ligand validation metrics for cryoEM data.

In conclusion, while our study demonstrates the potential of deep learning for ligand identification in cryoEM maps, it also highlights the need for continued research and development in this area. To foster further development in this field, we have made our training and testing data publicly available. We encourage the structural biology community to use and build upon these resources as addressing the challenges of map standardization and ligand validation will be crucial for realizing the full potential of cryoEM in drug discovery.

## Acknowledgements

The authors would like to thank the CCPEM mailing list for sharing their opinions on the causes of fundamental differences in the appearance of density histograms in EM Coulomb potential maps and X-ray electron density maps.

