## Supplement for "Ligand Identification in CryoEM and X-ray Maps Using Deep Learning"

### List of supplementary figures:

- **Fig S1**. Density value distributions for ten example X-ray and cryoEM difference maps.
- **Fig S2.** Tested deep learning architectures.
- **Fig S3.** Point sampling procedures.
- **Fig S4**. Heatmaps of model performance for different combinations of resolution and RSCC/Q-score.
- **Fig S5**. Heatmaps of model performance for different combinations of resolution and ligand size.
- **Fig S6**. Top-10 accuracy of each ligand group for different models on the X-ray ligand dataset.
- **Fig S7**. Top-10 accuracy of each ligand group for different MinkLoc3Dv2 training scenarios on the CryoEM ligand dataset.
- **Fig S8**. Top-10 accuracy of each ligand group for MinkLoc3Dv2 on the Mixed ligand dataset.
- **Fig S9**. Cumulative distributions of ligand groups for the X-ray, CryoEM, and Mixed test sets.
- **Fig S10**. Model performance based on the number of voxels in a ligand voxel grid.
- **Fig S11**. Model calibration plots for the X-ray, CryoEM, and Mixed test sets.

### List of supplementary tables:

- **Table S1**. Tested hyperparameter for the deep learning models.

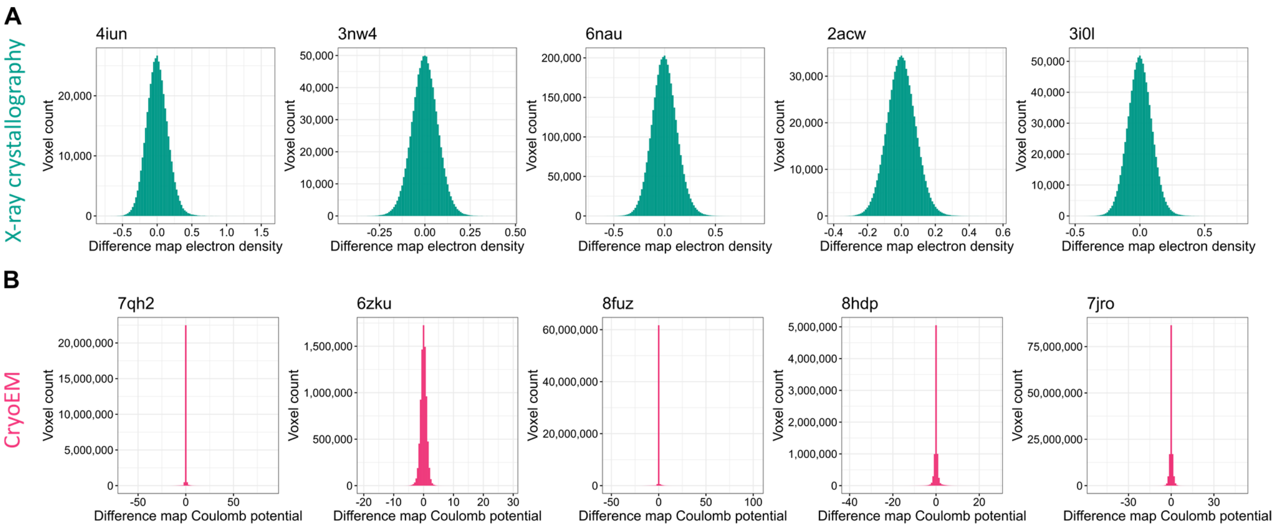


**Fig. S1** Density (voxel) value distributions for ten example X-ray and cryoEM difference maps. (**A**) Density value distributions for X-ray crystallography difference maps of PDB deposits 4iun, 3nw4, 6nau, 2acw, and 3i0l. (**B**) Density value distributions for cryoEM difference maps of PDB deposits 7qh2, 6zku, 8fuz, 8hdp, and 7jro. X-ray difference maps always have a Gaussian-like distribution with a slightly longer tail for the positive values. CryoEM difference maps usually have sharp spikes around zero and are much less consistent with each other. The selected PDB deposits correspond to the example ligands shown in Fig. 4 in the paper's main text.

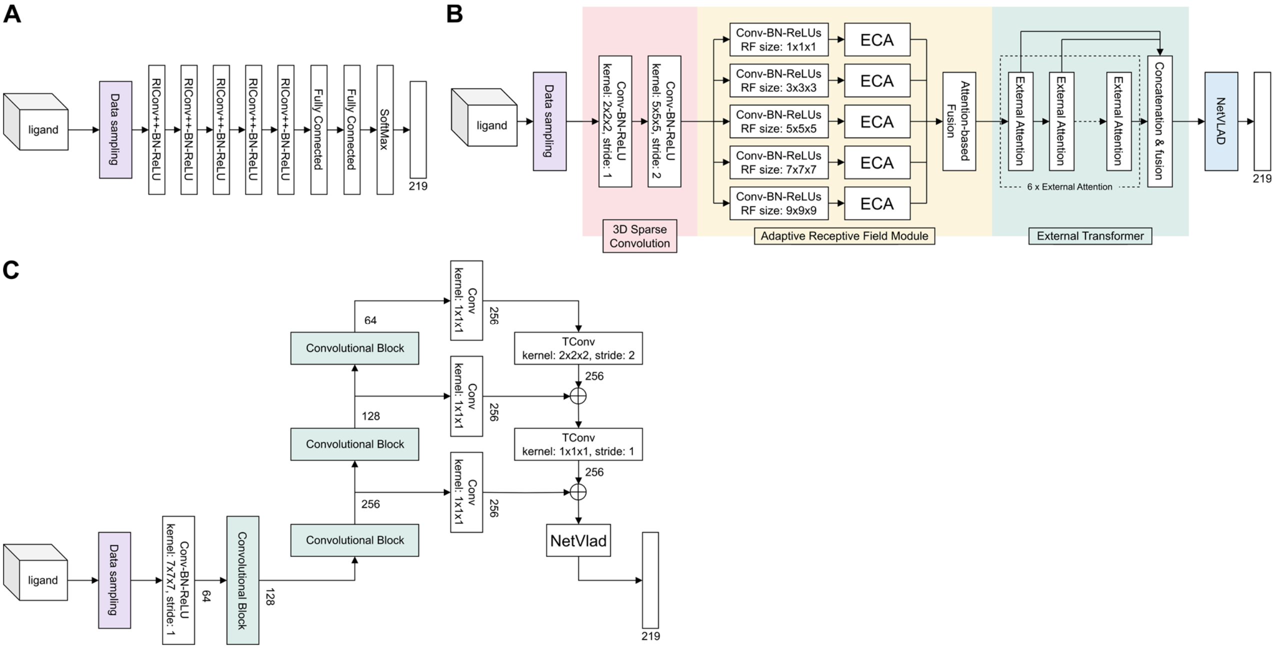


**Fig. S2** Schematics of deep learning architectures used to predict ligands. (A) The RiConv++ architecture has five enhanced rotation invariant convolution (RIConv++) layers.(B) The TransLoc3D architecture is built from four modules: 3DSparse Convolution, Adaptive Receptive Field, ExternalTransformer, and NetVLAD. (C) The MinkLoc3Dv2architecture utilizes information from a pyramid of three feature maps with different receptive fields. All the architectures were prepared to take as input the same sample of 2000 points and output the probability scores of all the studied 219 ligand groups.

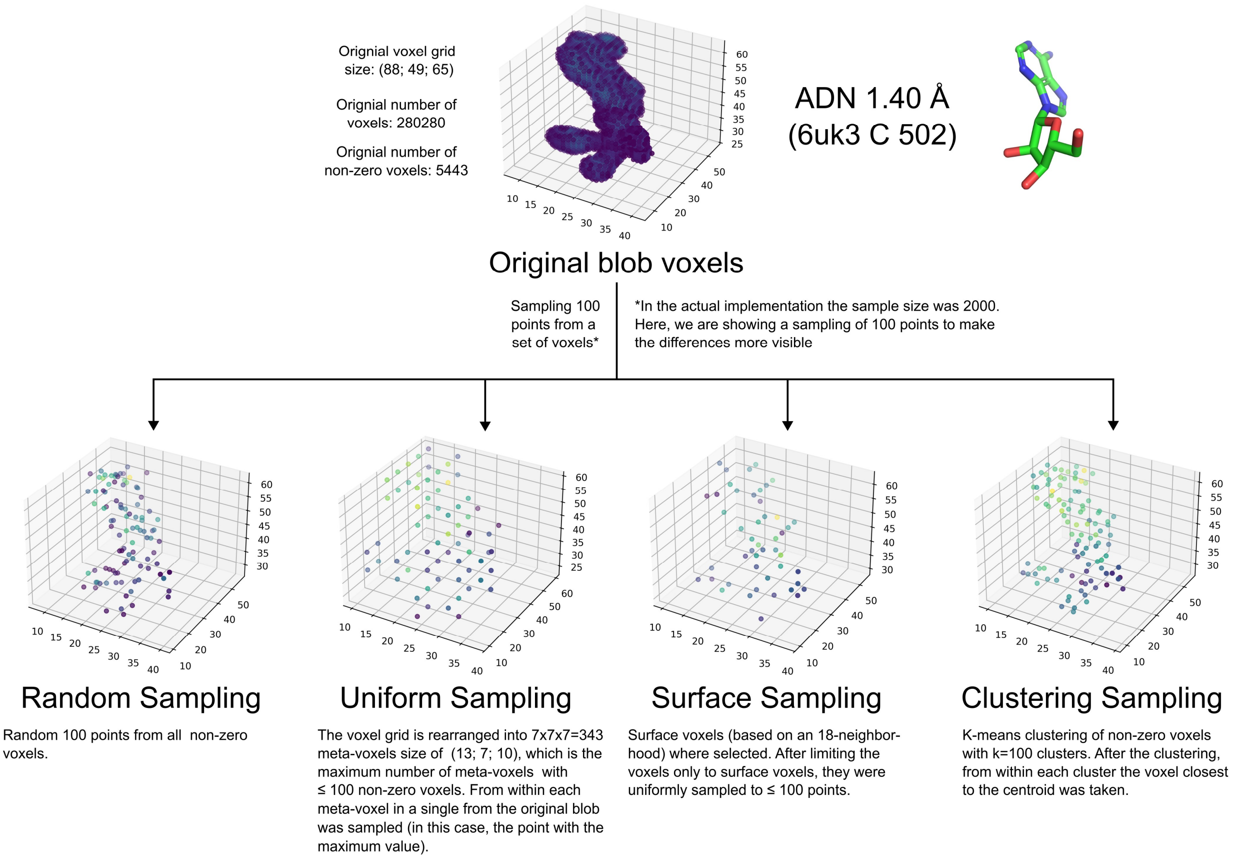


**Fig. S3** Voxel grid preprocessing approaches. In all the preprocessing approaches, the input is a full voxel grid (of any size) and the output is a set of at most $max_p$ points. For illustration purposes, the figure shows the effects of different sampling methods when $max_p$=100. In our deep learning models, $max_p$ was set to 2000.

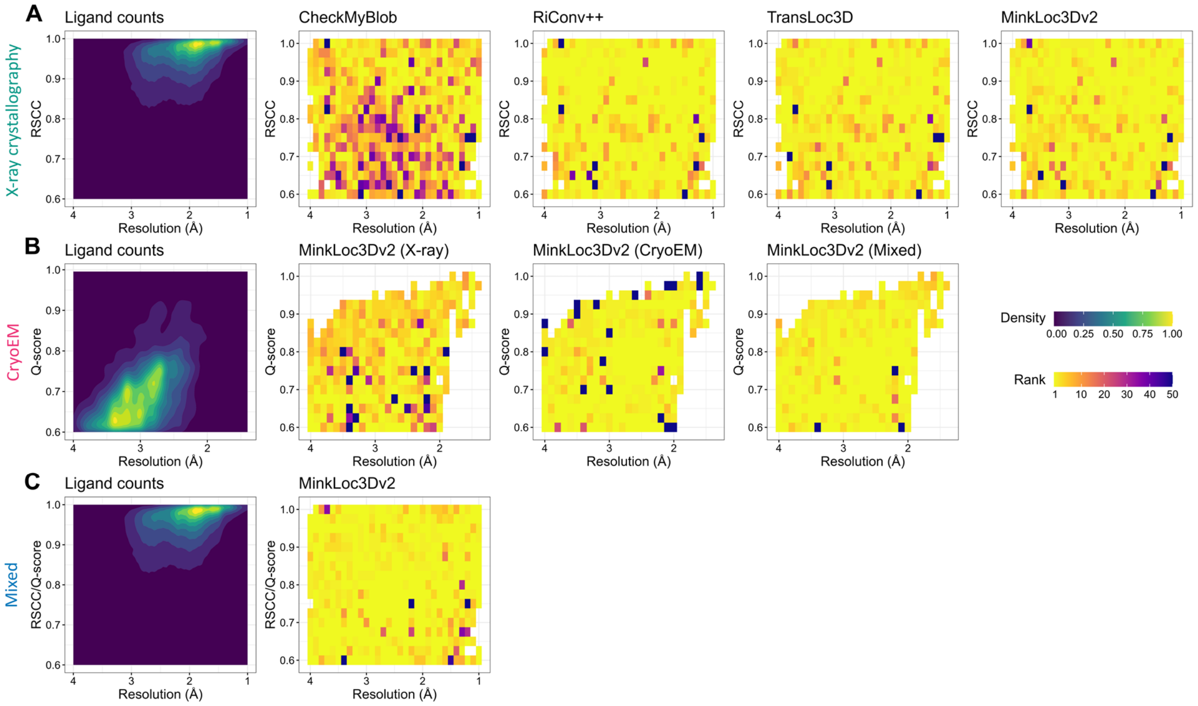


**Fig. S4.** Joint 2D ligand histograms and mean correct prediction rank heatmaps of resolution and real-space correlation coefficient (RSCC) for (**A**) X-ray, (**B**) cryoEM, and (**C**) mixed test datasets. Prediction ranks values above 50 capped at 50 to show the differences in top-ranking positions more clearly. It can be noticed that deep learning models (RiConv++, TransLoc3D, MinkLoc3Dv2) are better than CheckMyBlob in recognizing lower resolution and poor quality ligands on X-ray data (A). It can also be noticed how the combination of X-ray and cryoEM data complements each other when performing predictions on cryoEM (B) and mixed data (C).

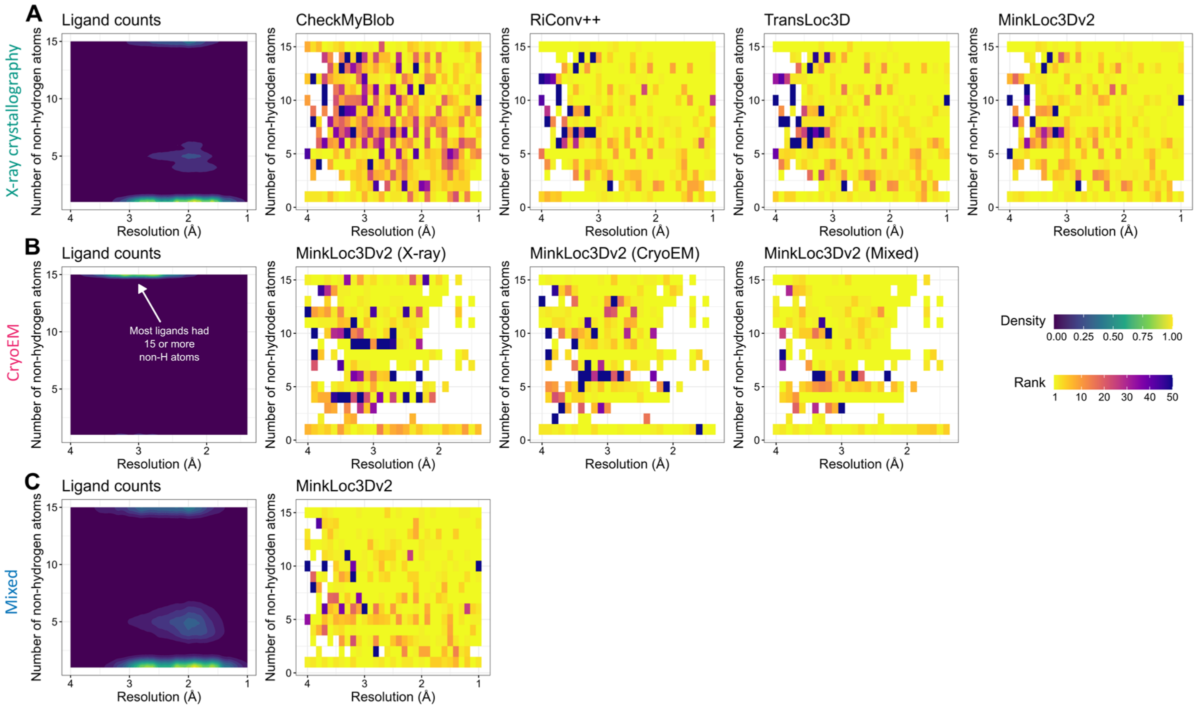


**Fig. S5.** Joint 2D ligand histograms and mean correct prediction rank heatmaps of resolution and real-space correlation coefficient (RSCC) for (**A**) X-ray, (**B**) cryoEM, and (**C**) mixed test datasets. Prediction ranks values above 50 capped at 50. The number of non-hydrogen atoms is capped at 15. It can be noticed that deep learning models (RiConv++, TransLoc3D, MinkLoc3Dv2) are better than CheckMyBlob in recognizing lower resolution and larger ligands on X-ray data (A). It can also be noticed how the combination of X-ray and cryoEM data complements each other when performing predictions on cryoEM (B) and mixed data (C).

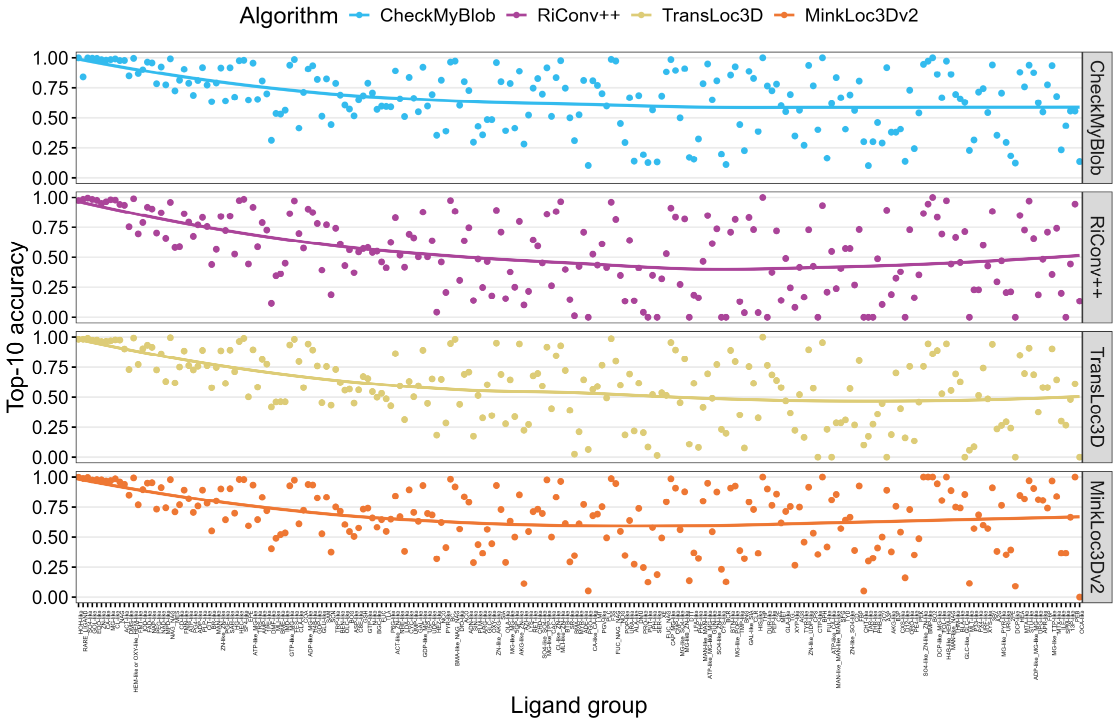


**Fig. S6.** Top-10 accuracy of each ligand group for different models on the X-ray ligand dataset. Ligands on the x-axis are sorted in descending order according to the number of examples in a given ligand group. Points represent top-10 accuracies for individual ligand groups; lines represent a spline regression fit to the sorted data.

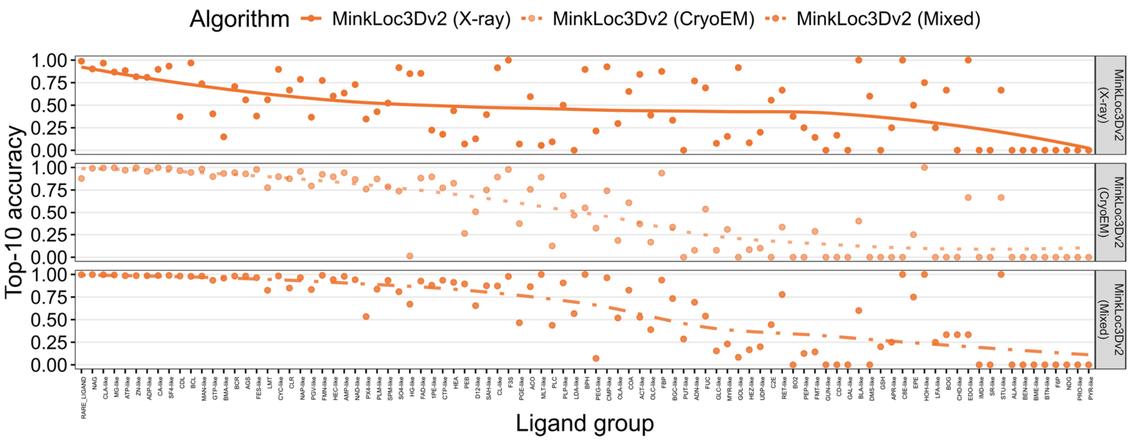


**Fig. S7.** Top-10 accuracy of each ligand group for different MinkLoc3Dv2 training scenarios on the CryoEM ligand dataset. Ligands on the x-axis are sorted in descending order according to the number of examples in a given ligand group. Points represent top-10 accuracies for individual ligand groups; lines represent spline regression fits to the sorted data.

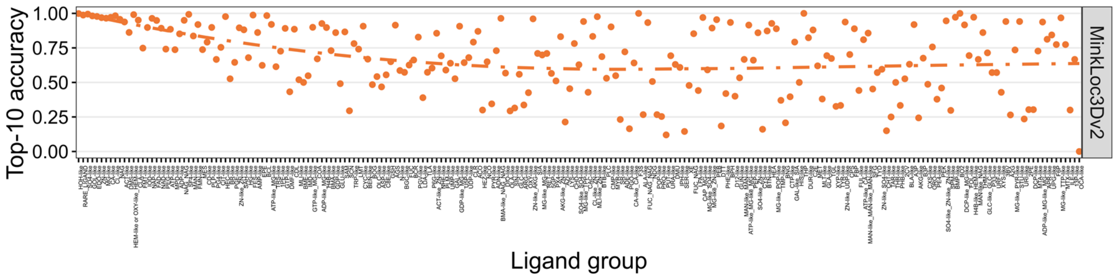


**Fig. S8.** Top-10 accuracy of each ligand group for MinkLoc3Dv2 on the Mixed ligand dataset. Ligands on the x-axis are sorted in descending order according to the number of examples in a given ligand group. Points represent top-10 accuracies for individual ligand groups; the line represents a spline regression fit to the sorted data.

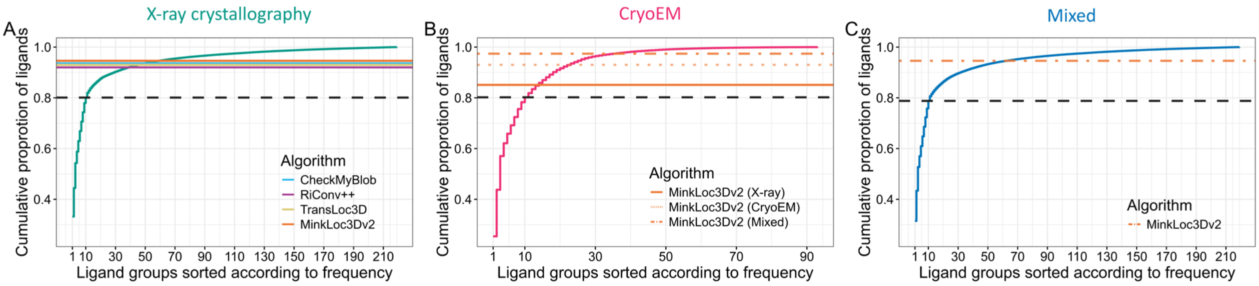


**Fig. S9.** Cumulative distribution of ligand groups for (**A**) X-ray crystallography test ligands, (**B**) cryoEM ligands, and (**C**) mixed dataset test ligands. Ligand groups are sorted and presented as numbers, with 1 corresponding to the most popular ligand group. Dashed black represents the proportion of the top 10 most popular ligand groups for a given dataset. Colored lines show the Top-10 Accuracy performance of the analyzed machine learning models.

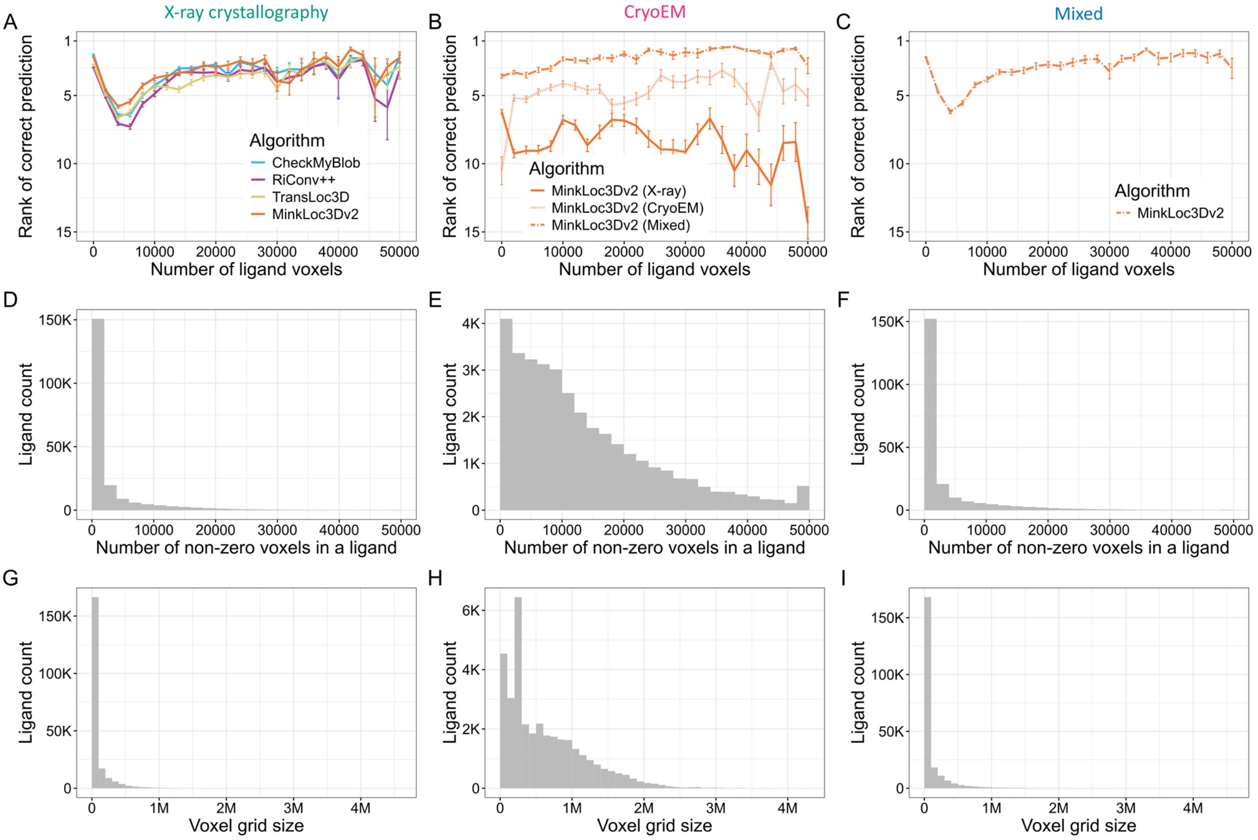


**Fig. S10.** Predictive performance of the analyzed machine learning for (**A**) X-ray crystallography, (**B**) cryoEM, and (**C**) mixed test datasets as a function of the number of (nonzero) ligand voxels on 0.2 Å voxel grid. Panels (**D**-**F**) show the corresponding ligand voxel counts whereas panels (**G**-**I**) show the corresponding voxel grid sizes (total number of voxels in a ligand grid) for X-ray crystallography, cryoEM, and mixed test data, respectively.

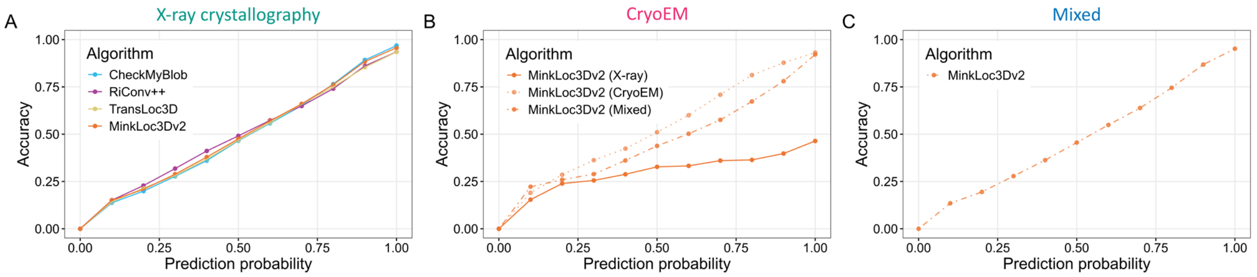


**Fig. S11.** Classification accuracy of the tested models within different prediction probability groups. (**A**) All the models are well-calibrated on the X-ray diffraction data. (**B**) On the cryoEM data, the model trained on X-ray ligands (MinkLoc3Dv2 (X-ray)) overestimates the higher prediction probabilities, whereas the cryoEM (MinkLoc3Dv2 (CryoEM)) and mixed training data (MinkLoc3Dv2 (Mixed)) are well-calibrated. (**C**) The mixed data model is also well-calibrated with the mixed testing data.

**Table S1**. Tested deep learning hyperparameters. The parameters that yielded the best performance on the validation set are highlighted in bold.

| Algorithm | Voxel sampling procedure | Batch size | Learning rate |
|---|---|---|---|
| RiConv++ | Random, Uniform, Surface, Clustering | 64 | 0.001 |
| RiConv++ | Random, Uniform, Surface, Clustering | 256 | 0.001 |
| RiConv++ | Random, Uniform, Surface, Clustering | 64 | 0.01 |
| RiConv++ | Random, Uniform, Surface, Clustering | 256 | 0.01 |
| TransLoc3D | Random, Uniform, Surface, Clustering | 64 | 0.001 |
| TransLoc3D | Random, Uniform, Surface, Clustering | 256 | 0.001 |
| TransLoc3D | Random, Uniform, Surface, Clustering | 64 | 0.01 |
| TransLoc3D | Random, Uniform, Surface, Clustering | 256 | 0.01 |
| **MinkLoc3Dv2** | Random, **Uniform**, Surface, Clustering | **128** | **0.001** |
| MinkLoc3Dv2 | Random, Uniform, Surface, Clustering | 196 | 0.001 |
| MinkLoc3Dv2 | Random, Uniform, Surface, Clustering | 256 | 0.001 |
| MinkLoc3Dv2 | Random, Uniform, Surface, Clustering | 512 | 0.001 |
| MinkLoc3Dv2 | Random, Uniform, Surface, Clustering | 784 | 0.001 |
| MinkLoc3Dv2 | Random, Uniform, Surface, Clustering | 128 | 0.01 |
| MinkLoc3Dv2 | Random, Uniform, Surface, Clustering | 784 | 0.01 |
